# Frequency modulations of cortical synchronization in human cortex during wakefulness and sleep

**DOI:** 10.64898/2026.03.13.710565

**Authors:** Maria Giovanna Canu, Gaia Burlando, Lorenzo Chiarella, Valentina Marazzotta, Marco Veneruso, Roberto Mai, Francesco Cardinale, Laura Tassi, Lino Nobili, Gabriele Arnulfo

## Abstract

Vigilance states are associated with reproducible reconfigurations of large-scale brain dynamics, reflected in changes in inter-areal phase synchronization and cross-frequency phase–amplitude coupling. Here, we characterized how these two forms of phase-based coordination jointly organize across sleep and wakefulness in the human brain at local and regional levels. We analysed phase-locking value (PLV) and phase–amplitude coupling (PAC) from intracranial stereo-electroencephalography (SEEG) recordings in 46 individuals with drug-resistant focal epilepsy, focusing on contacts outside the epileptogenic zone (non-epileptogenic zone, nEZ) to define physiological coupling profiles and comparing them with contacts within the EZ. For each subject, representative epochs of wakefulness, NREM sleep stages N2 and N3, and REM sleep were examined. Across vigilance states, large-scale phase synchronization exhibited distinct spectral fingerprints. Theta and sigma synchronization predominated during NREM sleep, beta synchronization increased during REM sleep, and theta interactions characterized wakefulness. PAC showed complementary state-dependent reorganizations: N3 was characterized by delta-driven modulation of broadband high-frequency activity; N2 additionally exhibited theta- and spindle-phase modulation of beta–gamma amplitudes; REM sleep showed reduced coupling; and wakefulness was marked by theta-to-beta interactions. Within vigilance states, epileptogenic regions displayed increased delta and gamma synchronization and enhanced delta-to-beta/gamma PAC, most prominently during N2 sleep and wakefulness, whereas differences between EZ and nEZ tissue were attenuated during REM sleep. Using partial least squares analysis, we further identified system-specific patterns of PLV–PAC covariation, with prominent involvement of temporal networks during NREM sleep and visual and limbic systems during REM sleep.

Together, these findings delineate a frequency-specific, state-dependent architecture linking phase synchronization and phase–amplitude coupling in the human brain and describe how epileptogenic networks deviate from physiological coupling profiles across vigilance states.

## Introduction

Neuronal oscillations support communication and information processing within and between brain regions (Bonnefond et al., 2017), enabling functional integration across multiple spatial and temporal scales (Fries, 2005; Cannon et al., 2014). Coordinated synchronization of brain activity reflects dynamic coupling between neuronal populations, indexing functional connections within neural circuits (Singer, 1999; Fries, 2005; Chiarion et al., 2023; Muñoz-Torres et al., 2023) and supporting the transient formation of functional networks (Palmigiano et al., 2017). By temporally aligning neuronal firing, synchronization enhances synaptic efficacy and promotes the selective routing of information across distributed brain systems, thereby enabling flexible reconfiguration of large-scale networks in response to internal and external demands (Fries, 2005; Palmigiano et al., 2017). Such oscillatory coordination contributes to attention, perception, and working memory (Fries, 2005; Engel and Fries, 2010), and has been proposed as a core mechanism for the large-scale integration of information (Tononi et al., 2016; Staresina et al., 2023).

Sleep provides a powerful framework for studying large-scale synchronization as unifying principles, as vigilance states modulate neuronal oscillations even in the absence of sensory input (Adamantidis et al., 2019).

Importantly, sleep is not homogeneous across the brain but shows heterogeneous spatial and temporal distributions (Vyazovskiy et al., 2011; Nobili et al., 2012; Geva-Sagiv and Nir, 2019).

Across sleep-wake cycles, neuronal synchrony varies even without external stimulation, indicating that intrinsic network dynamics govern the organization of inter-areal communication across vigilance states (Corsi-Cabrera et al., 1996, 2006).

To implement such large-scale coordination, neuronal communication relies on two complementary mechanisms: within-frequency synchronization and cross-frequency coupling (Varela et al., 2001; Fries, 2005; Canolty and Knight, 2010). Phase synchronization aligns extracellular activity at the same frequency across distant brain regions, enabling large-scale functional connectivity (Varela et al., 2001) even at fast rhythms (Arnulfo et al., 2020). In parallel, phase-amplitude coupling (PAC) integrates oscillations across spatiotemporal scales by modulating faster activity according to the phase of slower rhythms, including interactions between structures (Canolty and Knight, 2010). During NREM sleep, slow oscillations, spindles, and hippocampal ripples become tightly coordinated, creating a structured temporal framework for inter-areal communication (Cox et al., 2020; Staresina et al., 2023; Schreiner et al., 2024). Slow oscillations provide a global timing signal that gates the occurrence of spindles and ripples, thereby promoting local synchrony and long-range hippocampal–neocortical interactions. This nested organization opens windows during which memory traces can be reactivated and redistributed across cortical networks, a process that predicts subsequent memory consolidation (Geva-Sagiv et al., 2023).

In epilepsy, abnormal phase synchronization and excessive cross-frequency coupling characterize epileptogenic networks, reflecting hypersynchronous and aberrantly coupled neural activity (Lehnertz et al., 2009; Schmidt et al., 2014; Amiri et al., 2016). Sleep and epilepsy, indeed, share partially overlapping regulatory mechanisms of excitability and synchronization, such that oscillations supporting physiological coordination during sleep may, under pathological conditions, facilitate hypersynchronous discharges (Frauscher et al., 2015a; Roliz and Kothare, 2022; Sheybani et al., 2025). Consistently, PAC is enhanced both during deep sleep and within epileptogenic tissue (Amiri et al., 2016).

Building on this framework, the present study investigates phase synchronization, phase-amplitude coupling, and their interactions across vigilance states using human stereo-electroencephalography. We first characterize physiological coupling profiles in non-epileptogenic tissue and then quantify deviations in epileptogenic regions (Fig. 1A). By jointly quantifying phase-locking value (PLV) and PAC within a large-scale network, we identify frequency-specific patterns of inter-areal communication across sleep stages and assess their role in organizing brain-wide dynamics during NREM and REM sleep (Fig. 1B).

**Figure 1:**
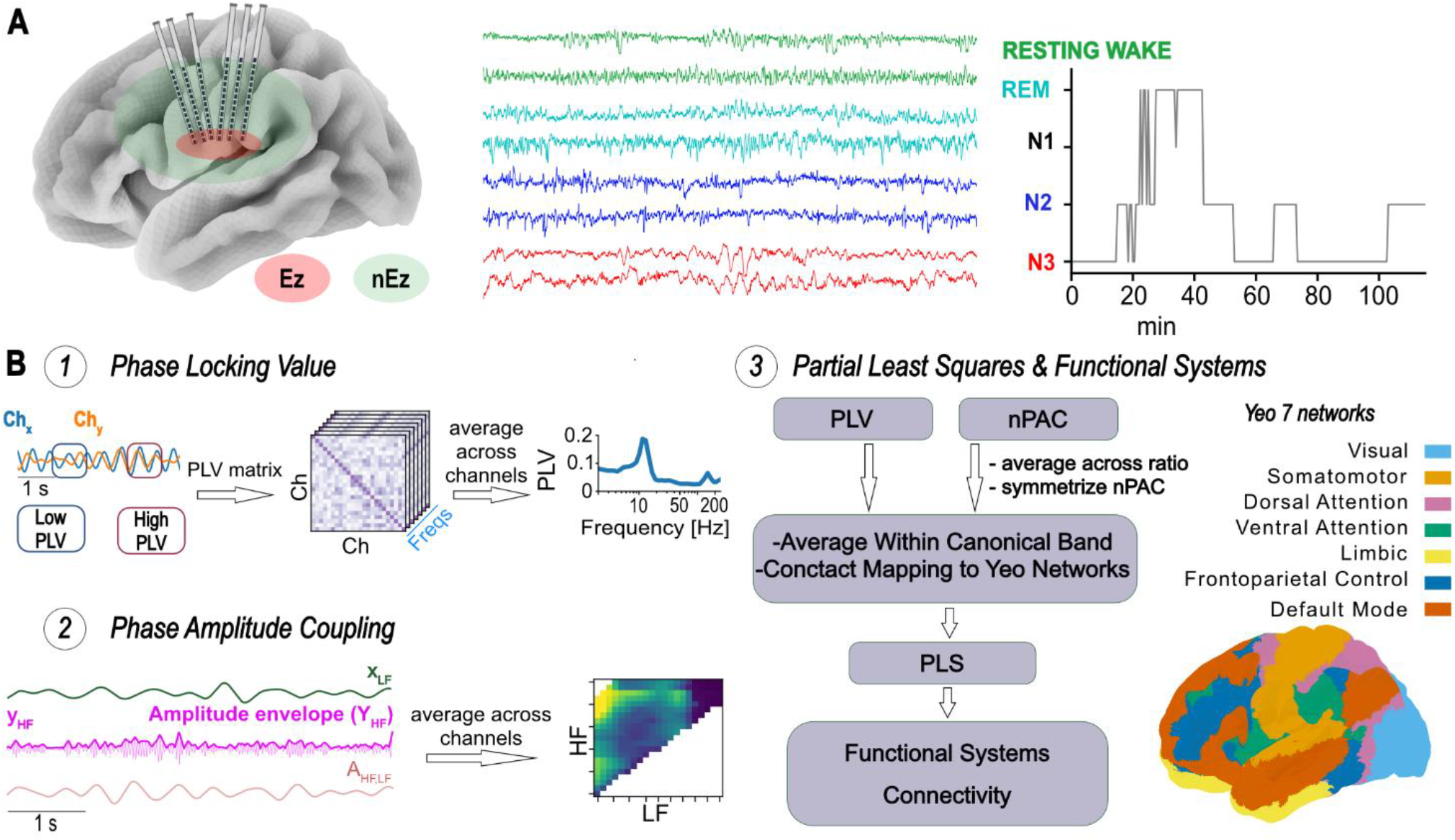
Study schematic and analysis workflow. **(A)** Schematic representation of the experimental setup and sleep staging procedure. Intracranial recordings were obtained using depth electrodes implanted in epileptogenic (EZ) and non-epileptogenic (nEZ) regions. Representative signal traces are shown for resting wakefulness and sleep stages (REM, N2, N3), together with an example hypnogram illustrating vigilance-state segmentation across the recording period. Only contacts located in non-epileptogenic zones were included in the main analyses, while epileptogenic zones were considered in comparative analyses. **(B)** Connectivity analysis workflow. (1) Band-pass filtered signals from two channels are used to estimate phase synchrony across frequencies. For each frequency band, PLV is computed between all channel pairs, yielding channel × channel connectivity matrices. PLV values are then averaged across channel pairs to obtain a frequency-resolved PLV profile. (2) PAC quantifies the interaction between the phase of low-frequency (LF) oscillations and the amplitude envelope of high-frequency (HF) activity. The HF signal is transformed into its amplitude envelope and filtered at LF to extract its slow modulation. PAC is then computed between LF phase and HF amplitude across frequencies, yielding LF × HF coupling matrices that are averaged across channel pairs. (3) PLV and normalized PAC (nPAC) measures are averaged within canonical frequency bands and mapped from contact-level connectivity to cortical regions belonging to the Yeo 7 functional networks. nPAC values are first averaged across high-frequency ratios and symmetrized. Partial least squares analysis is then applied to identify shared modes of connectivity between PLV and nPAC. The resulting components are used to reconstruct connectivity patterns between large-scale functional systems.

## Methods

### Data acquisition

We examined stereo-electroencephalography (SEEG) data from 41 individuals diagnosed with drug-resistant focal epilepsy (DRE) who underwent pre-surgical clinical evaluation to identify the epileptogenic focus for potential ablation. A schematic illustration of the SEEG electrode implantation is shown in Fig. 1A (left panel). Before electrode implantation, the participants provided written informed consent to participate in research studies. This study received approval from the Niguarda Ca’ Granda Hospital, Milan’s ethical committee (ID 939), and it follows the principles outlined in the Declaration of Helsinki. We obtained monopolar local-field potentials (LFPs) from brain tissue using multi-lead platinum-iridium electrodes. Each penetrating shaft featured 8 to 15 contacts, measuring 2 mm in length, 0.8 mm in thickness, and an inter-contact distance of 1.5 mm (manufactured by DIXI medical, Besancon, France). We acquired around 10 minutes of continuous spontaneous brain activity from these patients with eyes closed (resting state) and one overnight recording (7.4 ± 0.9 h) of SEEG and polysomnography, including EOG, EMG, and scalp EEG electrodes the latter located at Fz, Cz, Pz, C3, P3, C4, and P4.

### Sleep scoring

Sleep specialists manually scored the full-night recordings in 30-second epochs according to the American Academy of Sleep Medicine (AASM) criteria, using scalp EEG, EOG, and submental EMG. From these annotations, we extracted consecutive five-minute segments for each sleep sub-stage (N3, N2, REM) as well as for wakefulness. Continuous N2 and REM epochs were selected from the initial hours of sleep, whereas N1 stages were excluded due to their short duration relative to the other stages. For N3 sleep selection, slow-wave activity (SWA) was quantified by extracting the δ-band amplitude envelope. Signals were band-pass filtered (0.5–4 Hz) using a third-order Butterworth filter. To prevent phase distortion, the filter was applied in both forward and reverse directions (zero-phase filtering) using a second-order sections implementation. The instantaneous amplitude envelope was subsequently computed via the Hilbert transform. A five-minute segment centred on the recording’s maximum SWA peak was then extracted as the representative N3 epoch. In total, N3 and wake epochs were extracted from 41 subjects, N2 epochs from 31 subjects, and REM epochs from 26 subjects. Fig. 1A (right panel) shows a representative hypnogram together with example SEEG recordings from one subject.

### Signal pre-processing

We adopted the closest white-matter (cWM) referencing method, in which grey-matter contacts are referenced to the nearest white-matter contact. This approach minimizes interference from active sources and ensures consistent signal polarity, thereby improving the precision of phase estimates (Arnulfo et al., 2015). Line noise at 50 Hz and its harmonics up to the Nyquist frequency were removed using IIR notch filters. Each signal was then decomposed using a bank of 40 Morlet wavelet filters with central frequencies ranging from 2 to 250 Hz. Epileptic events, such as interictal spikes, are characterized by brief, high-amplitude transients with widespread spatial propagation. To prevent these events from artificially inflating synchrony measures, we excluded 500-ms windows containing interictal epileptic events (IIEs). IIEs were identified as periods in which at least 10% of cortical channels simultaneously exhibited sharp amplitude peaks, defined as envelope values exceeding five standard deviations above the mean in more than half of the 40 frequency bands.

For the first part of the analysis, only contacts located in non-epileptogenic zones (nEZ) were included to ensure that the results reflected physiological activity. In a subsequent analysis, phase locking value (PLV) and phase-amplitude coupling (PAC) were compared between contacts within epileptogenic zones (EZ) and those in nEZ to assess potential alterations in synchronization and coupling patterns.

### Connectivity & clustering

To investigate inter-areal phase synchronization, we used the Phase Locking Value (PLV). The PLV is computed as the absolute value of the complex PLV (cPLV), which is derived from the complex wavelet coefficients of the signals at a given frequency. Specifically, if x′(t)and y′(t) represent the complex wavelet coefficients of two signals, the cPLV is defined as:

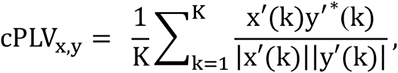

where K is the total number of samples and * denotes the complex conjugate. The PLV is then obtained as:

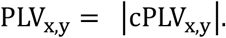

Additionally, we used the imaginary part of cPLV (iPLV), defined as iPLV = |Im(cPLV)|, a metric that is insensitive to zero-lag interactions attributed to volume conduction (Palva et al., 2018). Both metrics provide a scalar measure ranging from 0 (no synchronization) to 1 (perfect phase locking).

To identify patterns across vigilance states, we concatenated the four stage-specific PLV spectra for each subject and performed subject-level agglomerative hierarchical clustering using the Euclidean distance metric. The number of clusters was fixed at four.

### Phase-amplitude coupling

Phase-amplitude coupling (PAC) provides information about the correlation between the phase of slow oscillations and the amplitude of faster rhythms. We computed PAC between pairs of low-frequency (LF) and high-frequency (HF) components. If θ_x,LF_(*t*) denotes the phase of the LF signal from channel x and 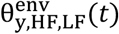 the phase of the amplitude envelope of the HF signal from channel y, filtered at LF, PAC was defined as:

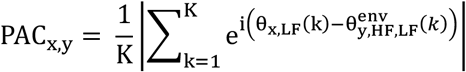

where K is the total number of samples. Similar to the PLV, the PAC also offers a measure in the 0–1 range, with 1 denoting full phase-amplitude interaction and 0 denoting no interaction. To account for spurious coupling, we used a normalized PAC (nPAC) defined as PAC_PLV,obs_/PAC_PLV,sur_, which represents PAC above the null hypothesis level. nPAC values were discarded when the corresponding HF component exceeded 200 Hz. Our analysis focused on inter-areal interactions, specifically assessing how the low-frequency activity of one channel modulated the high-frequency activity recorded from another channel.

PLV and PAC were computed using the Python toolbox CROCOpy for the evaluation of brain criticality and connectivity (Myrov et al., 2026).

### Neuroanatomical organization

We identified the anatomical location of each recording contact in individualised pre-surgical MRI using the SEEG-Assistant module (Narizzano et al., 2017). This tool segments the position of individual contacts visible in post-implant CT scans to pre-implant MRI, enabling the accurate identification of each contact position with respect to the patient’s brain anatomy. Contact locations were then mapped onto a standard functional brain atlas. We utilized the Schaefer parcellation with a resolution of 200 parcels (Schaefer et al., 2018), which were generated based on individual pre-surgical 3D T1-weighted (FFE) MRI scans and processed using the Freesurfer software (Fischl, 2012). Finally, we combined parcels into 7 functional systems (Yeo et al., 2011) using the parcel-to-system mapping provided in the Schaefer atlas.

### Partial least squares (PLS)

Partial least squares (PLS) is a statistical method that finds linear combinations of predictor variables X and response variables Y that maximize the covariance between them, focusing on predicting Y from X.

Given two datasets X (size n × p) and Y (size n × q), PLS finds weight vectors w and c such that the projections t = Xw and u = Yc maximize the covariance cov(t, u).The algorithm iteratively extracts latent components by solving the following optimization:

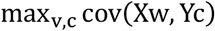

After computing the scores t and u, the datasets are deflated to remove the information captured by the current component, and the procedure is repeated for the next component. The resulting latent variables summarize the most predictive information in X for Y, and the weights w and c can be used to interpret the contribution of the original variables.

### Preprocessing of connectivity matrices

PLS was applied between nPLV and nPAC to identify shared modes of connectivity. Because nPAC is computed over low-frequency phase × high-frequency amplitude, we collapsed the high-frequency dimension by averaging across ratios (HF amplitude ~30–200 Hz; >200 Hz excluded). This yields one PAC value per low-frequency phase band (δ: 2–4 Hz, θ: 4–8 Hz, α: 8–12 Hz, β: 12–30 Hz), reflecting modulation of broadband HF activity rather than HF-band-specific coupling. The contribution phase of the γ band was not considered, as no phase-amplitude coupling was observed in modulating faster rhythms (Fig. 3). Edges corresponding to functional systems not sampled by SEEG in each subject were excluded from further analyses. For each vigilance state, PLS was applied using bands as features to identify linear combinations of nPLV and nPAC that maximally covaried. The first latent component was extracted, and the scores were reconstructed into the original connectivity matrices for each subject. The number of subjects included varied across vigilance states (N3: 38, N2: 27, REM: 23, wake: 38). Finally, we computed the average connectivity across X and Y scores and then across subjects to obtain group-level representations of the PLS-derived components.

### Null hypothesis and statistic

We computed surrogate cPLV (PLV and iPLV) and surrogate PAC for every channel pair to determine the null hypothesis distributions for our measures. We generated surrogate data that eliminated correlations between two contacts while preserving the temporal autocorrelation structure of the original signals. We created the surrogate by dividing each narrow-band time series for each contact pair into two blocks with a random time point k:

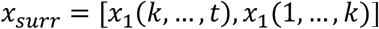

Surrogate analysis was applied with two distinct purposes, depending on the metric. For PLV, surrogate levels were incorporated into the main phase synchronization spectra to provide a visual reference for the noise level in the data. For PAC, surrogate analysis was used to normalize PAC values, as detailed in the corresponding analysis section (see: Phase–amplitude coupling).

For within-subject pairwise comparisons between vigilance states, we employed the Wilcoxon signed-rank test for both PLV and PAC measures. Comparisons across all four vigilance states were performed using the Kruskal–Wallis test. In contrast, Wilcoxon signed-rank tests were used to compare PLV and PAC between EZ and nEZ. For all statistical analyses, p-values were corrected for multiple comparisons across frequencies using the Benjamini–Hochberg procedure (α = 0.05) (Genovese et al., 2002). In addition, effect sizes were computed for all conditions to quantify the magnitude of differences independently of sample size. Effect sizes were estimated using Cohen’s *d* for pairwise comparisons and η^2^ for multiple-group comparisons.

## Results

### Phase synchronisation spectra across different behavioural states

In this work, we assessed synchrony of neuronal oscillations across different vigilance states. We chose the Phase-Locking Value (PLV) to assess the inter-areal phase synchronization between all nEZ contact pairs. PLV decayed as a function of frequency and showed multiple synchronisation peaks (Fig. 2A) that cannot be attributed to spurious synchronisation due to volume-conduction (Fig. S1). Comparing PLV across all vigilance states, we identified two main frequency ranges where significant (*p* < 0.05, Kruskal–Wallis test, Benjamini– Hochberg (BH) correction, α = 5%, Fig. 2B) differences emerged: from the θ band up to 15 Hz (0.01 < η^2^ < 0.06, medium effect size and η^2^ > 0.14 large effect size; respectively), and within the low β range (20–30 Hz; 0.01 < η^2^ < 0.06, medium effect size). Post-hoc pairwise assessment (Wilcoxon signed-rank test with BH correction for multiple comparisons, α = 0.05) revealed that θ oscillations (5–10 Hz) exhibited significantly higher synchronization during both wakefulness and NREM sleep compared to REM sleep (*p*<0.05; |d|> 0.8 large effect size; Fig. S2). In the spindle band (12–15 Hz), PLV showed a pronounced peak during N3 sleep, a secondary peak during N2, and significantly lower synchronization during both wakefulness and REM sleep (*p* < 0.05 for REM vs. N3, wake vs. N3, and REM vs. N2; |d|> 0.8 large effect size; Fig. S2). In the β band (15–30 Hz), PLV was highest during REM sleep, with a significant increase compared to N3 (*p* < 0.05; |d|> 0.8 large effect size; Fig. S2). Moreover, in the ripple band (70–100 Hz), post-hoc pairwise comparisons revealed significantly greater synchronization during REM sleep compared to wakefulness (*p* < 0.05; |d|> 0.8 large effect size; Fig. S2).

**Figure 2:**
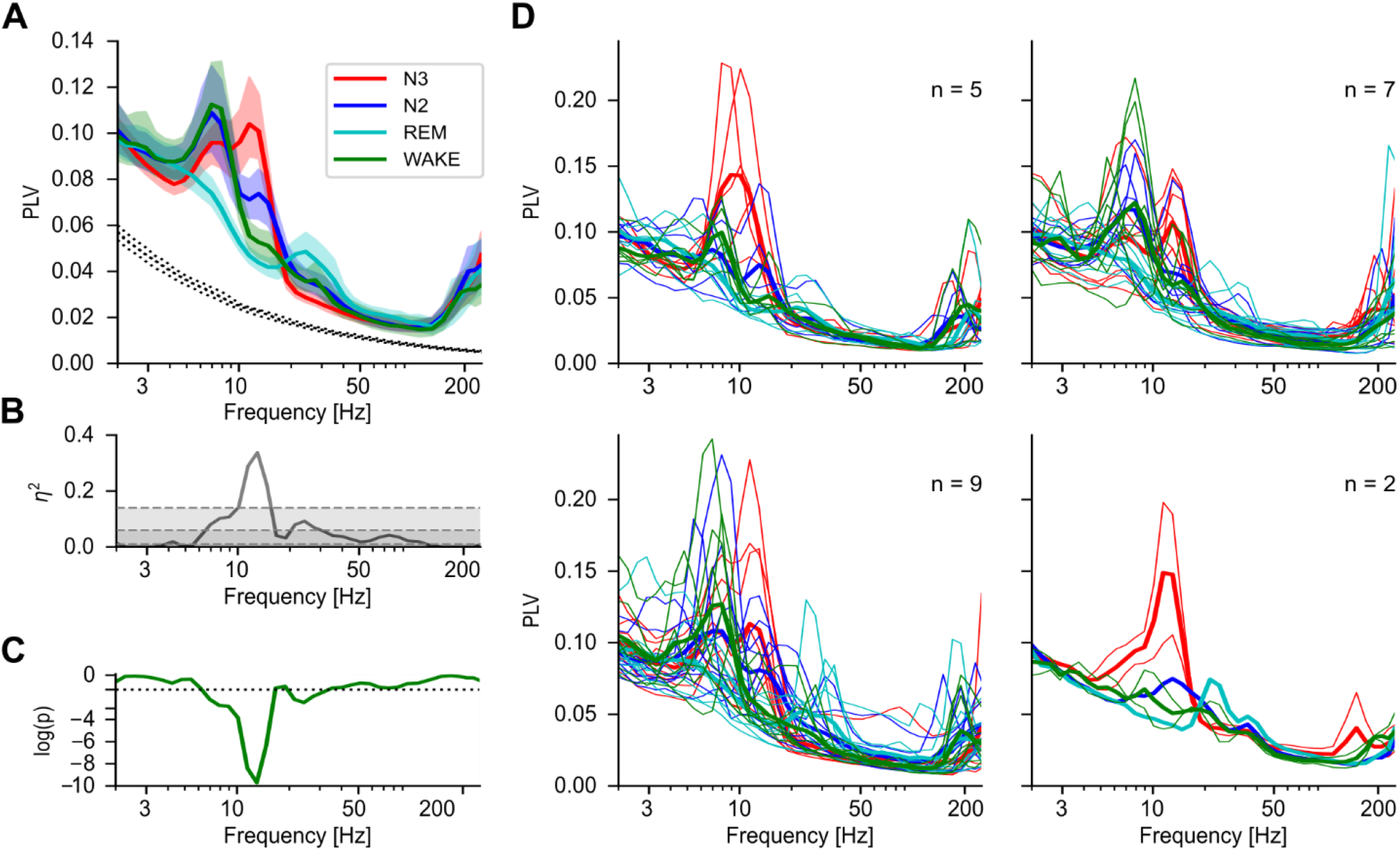
Phase synchronization across vigilance states and spectral clustering of individual PLV profiles. **(A)** Phase synchronization spectra for N3 (red), N2 (blue), REM (cyan), and wake (green) conditions across subjects in nEZ. Phase synchronization was computed as the phase locking value. The shaded areas represent the 2.5th to 97.*5th percentile bootstrap (N=1000) confidence limits around the mean (thick lines)*. **(B)** Effect size computed with eta squared (η^2^) for the comparison of PLV between the four vigilance states. Coloured areas represent the value range for small (η^2^ < 0.01, dark grey), medium (0.01 < η^2^ < 0.06, light grey), and large (η^2^ > 0.14, no shadow) effect sizes. **(C)** p-values profiles for the comparison of PLV between the four vigilance states (Kruskal–Wallis test, Benjamini–Hochberg correction, α = 0.05). The black dashed line denotes the significance threshold (p = 0.05). **(D)** Clustering of individual PLV spectra using a multi-feature approach. Thick lines represent the mean PLV of each cluster, while thinner lines show the spectra of individual subjects within each cluster.

**Figure 3:**
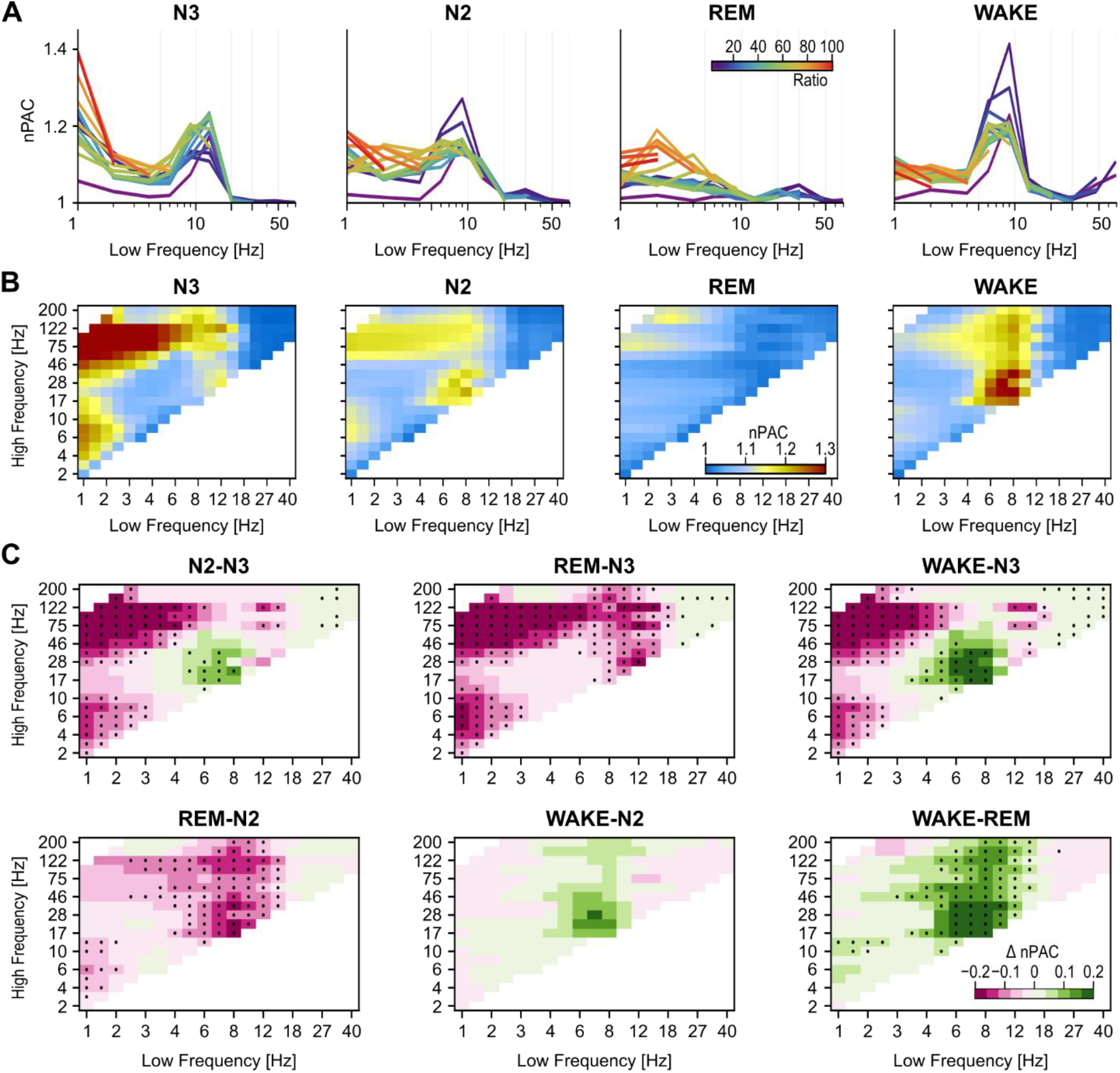
Normalized phase–amplitude coupling (nPAC) across frequencies and vigilance states. **(A)** Individual nPAC spectra, where each coloured line represents a specific phase–amplitude frequency pair (PAC ratio), illustrating how coupling strength varies with the low-frequency phase (x-axis, logarithmic scale), while the y-axis reflects normalized PAC values. **(B)** nPAC colormaps showing the modulation between low-frequency phase (x-axis) and high-frequency amplitude (y-axis). **(C)** Differences in nPAC between vigilance states, with dotted points marking frequency pairs with significant differences p < 0.05; Wilcoxon rank-sum test, Benjamini–Hochberg correction, α = 0.05, and |d| > 0.2; Cohen’s d effect size.

Furthermore, we explored the inter-individual variability in synchronization profiles (Fig. 1D). We evaluated the similarity of individual PLV spectra in the 23 subjects that were consistently recorded in all four stages. The subjects clustered into distinct groups: the first cluster (N=5 subjects) exhibited PLV spectrum during N3 characterized by a single peak in the θ-σ band, whereas N2 displayed two similar peaks in the θ and σ bands. The second cluster (N=7 subjects) featured different peaks in N3 within the θ and σ bands, but lacked two distinct peaks for wakefulness and N2 in the σ band, a characteristic observed in the third cluster (N=9 subjects), where N2 shows a high synchronisation peak in the σ band. In the third cluster, REM sleep shows a peak in the β band. Finally, the last cluster (only two subjects) presented a flattened profile of PLV, except for the high peak in the θ-σ band for N3.

### Phase-amplitude coupling across vigilance states

Normalized PAC (nPAC) exhibited clear stage-dependent profiles (Fig. 3). In N3, coupling was strongest for δ-phase oscillations (~1–3 Hz)—consistent across all ratios (Fig. 3A), indicating robust slow-oscillation nesting of higher-frequency activity (Fig. 3B). N3 also showed a secondary dominant interaction in the spindle range (10–15 Hz), influencing broadband high frequencies (~30–200 Hz) (Fig. 3B). In N2, δ-phase–driven coupling remained prominent and broadly distributed across high frequencies. By contrast, dominant θ (5–10 Hz) and spindle (10–15 Hz) phase interactions were more selectively associated with β/low-γ amplitudes (~30–100 Hz) (Fig. 3A–B). During REM, nPAC was globally attenuated across low-frequency phases, with an increase in the δ range (1–4 Hz) at high phase-to-amplitude ratios—corresponding to high-γ/ripple frequencies. In wakefulness, the dominant coupling shifted to a θ-centred maximum (5–10 Hz) that spanned a wide range of ratios but was particularly pronounced in the β band (~15–30 Hz).

The pairwise difference maps delineate state-specific nPAC patterns across vigilance states (Fig. 3C), obtained using the Wilcoxon signed-rank test with Benjamini–Hochberg correction for multiple comparisons (α = 0.05). In the N3–N2 contrast, N3 showed higher δ-phase (1–3 Hz) coupling with high-frequency activity amplitude, whereas N2 presented increased spindle-phase (10–15 Hz) interactions with β/low-γ rhythms amplitudes (30– 100 Hz). Relative to REM, N3 displays coupling across the entire slow-to-γ/ripple range. In the N3– wakefulness comparison, slow-oscillation nesting is observed in N3, while wakefulness exhibits α (8–12 Hz) to β/low-γ coupling.

In the REM–N2 contrast, N2 showed PAC involving spindle-phase activity (10–15 Hz) with β/low-γ amplitudes (30–100 Hz), while REM displays a general reduction of coupling. In the Wake–N2 comparison, PAC patterns within the examined frequency ranges were comparable to those observed during wakefulness, and no statistically significant differences were detected. In the Wake–REM contrast, PAC during wakefulness extends across θ and α and high-frequency amplitudes, while REM shows reduced engagement across this range.

### Phase synchronization & phase-amplitude coupling in the epileptogenic zone

We next tested whether recordings from the epileptogenic zone (EZ) exhibited altered synchronization relative to non-epileptogenic pairs. We computed population-average phase-locking value (PLV) and phase-amplitude coupling (PAC) for EZ–EZ pairs and compared these to nEZ–nEZ pairs, applying Wilcoxon signed-rank with Benjamini–Hochberg correction for multiple comparisons. Overall, EZ pairs showed modest increases in both PLV and PAC across vigilance states (Fig. 4).

**Figure 4:**
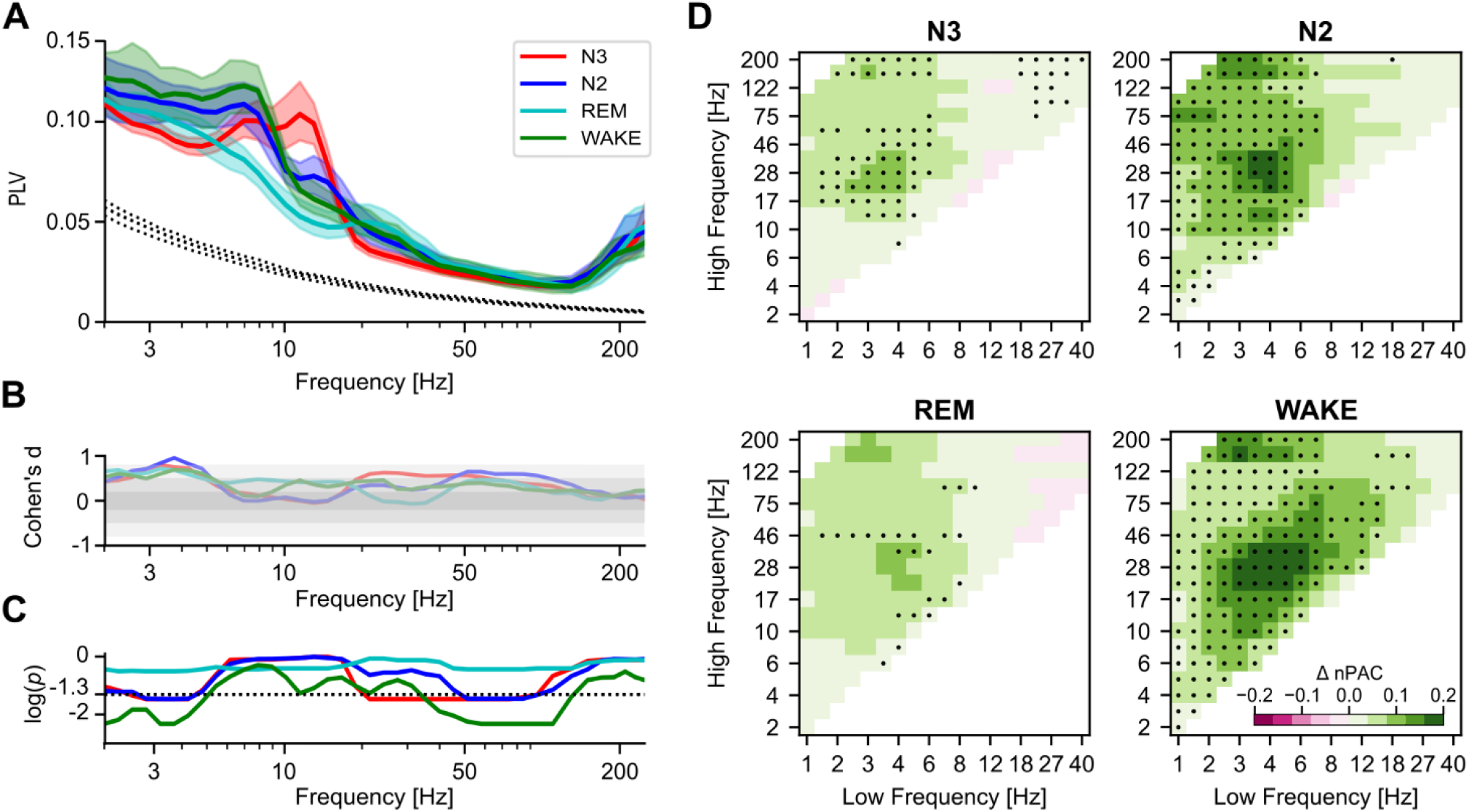
Effect of epilepsy on the phase locking value and PAC within conditions. **(A)** PLV spectrum for the PLV of EZ-EZ channel pairs averaged across subjects for the three vigilance states, including N3 (red), N2 (blue), REM (cyan), and Wake (Green). Thick lines represent population average, while shaded areas represent the 95% bootstrapped confidence intervals around the mean (N=1’000). Dashed lines represent the noise level computed with surrogates (N=100). **(B)** Effect size computed with Cohen’s d for the comparison of PLV between EZ-EZ and nEZ-nEZ channel pairs for individual vigilance states. Coloured areas represent the value range for small (|d|<0.2, dark grey), medium (0.2<|d|<0.8, light grey), and large (|d|> 0.8, dark grey) effect sizes. **(C)** p-values from the Wilcoxon signed-rank test for the comparison of PLV between EZ-EZ and nEZ-nEZ channel pairs for individual vigilance states. The black dashed line denotes the significance threshold (p = 0.05). **(D)** PAC comodulogram of the differences between EZ-EZ and nEZ-nEZ PAC. Dotted points mark frequency pairs with significant differences (p < 0.05 and |d| > 0.2).

Phase synchronization values showed increased values in the δ and γ bands (Fig. 3A), more pronounced in the slow oscillations range with a strong effect size (d>0.8) for N2, N3, and wake conditions, while γ-band activity only showed a small to medium effect (Wilcoxon signed-rank, *p* < 0.05, BH corrected). Phase-amplitude coupling showed a similar trend to PLV, with overall δ-to-β up to γ and high-γ being the most affected frequency modes (Fig. 4B) with only a small effect size (d < 0.2), highest in N2 and wake conditions. Overall, these results suggest that within and cross-frequency coupling between channel pairs recording from the epileptogenic zone are increased in δ and γ and in δ-to-γ coupling with a medium effect.

### Joint synchrony between functional systems exhibits frequency-specific patterns

To investigate which functional systems are most synchronized, we applied the PLS to relate large-scale phase synchrony with cross-frequency coupling. The subject-average scores revealed that the patterns of synchrony were vigilance-state-dependent, with distinct functional systems emerging as more or less engaged across vigilance states. In particular, in NREM stages (*i*.*e*., N2 and N3), the temporal system (Temp) was the node with the highest degree (s_N2,Temp_ = 21.63, s_N3,Temp_ = 25.72) (Fig. 5A–B). In N2, the highest contribution was from θ PAC (w = 28.41%), whereas δ frequencies were most prominent in N3 (w = 27.35%) (Fig. 5C). During the resting state, no single system clearly dominated; the peripheral visual system (VisP) showed higher node strength (s_rest,VisP_ = 20.16), with PLV weights concentrated in the α band (w = 27.16%) and PAC weights in the β band (w = 27.52%). In REM sleep, the limbic (Limb) and peripheral visual systems (VisP) exhibited the highest node strength (s_REM,Limb_ = 51.06 and s_REM,VisP_ = 50.96, respectively). δ-PLV and β-PAC contributed the most in the PLS model (w = 28.66% and w = 34.43%, respectively), capturing the covariation between PLV and PAC.

**Figure 5:**
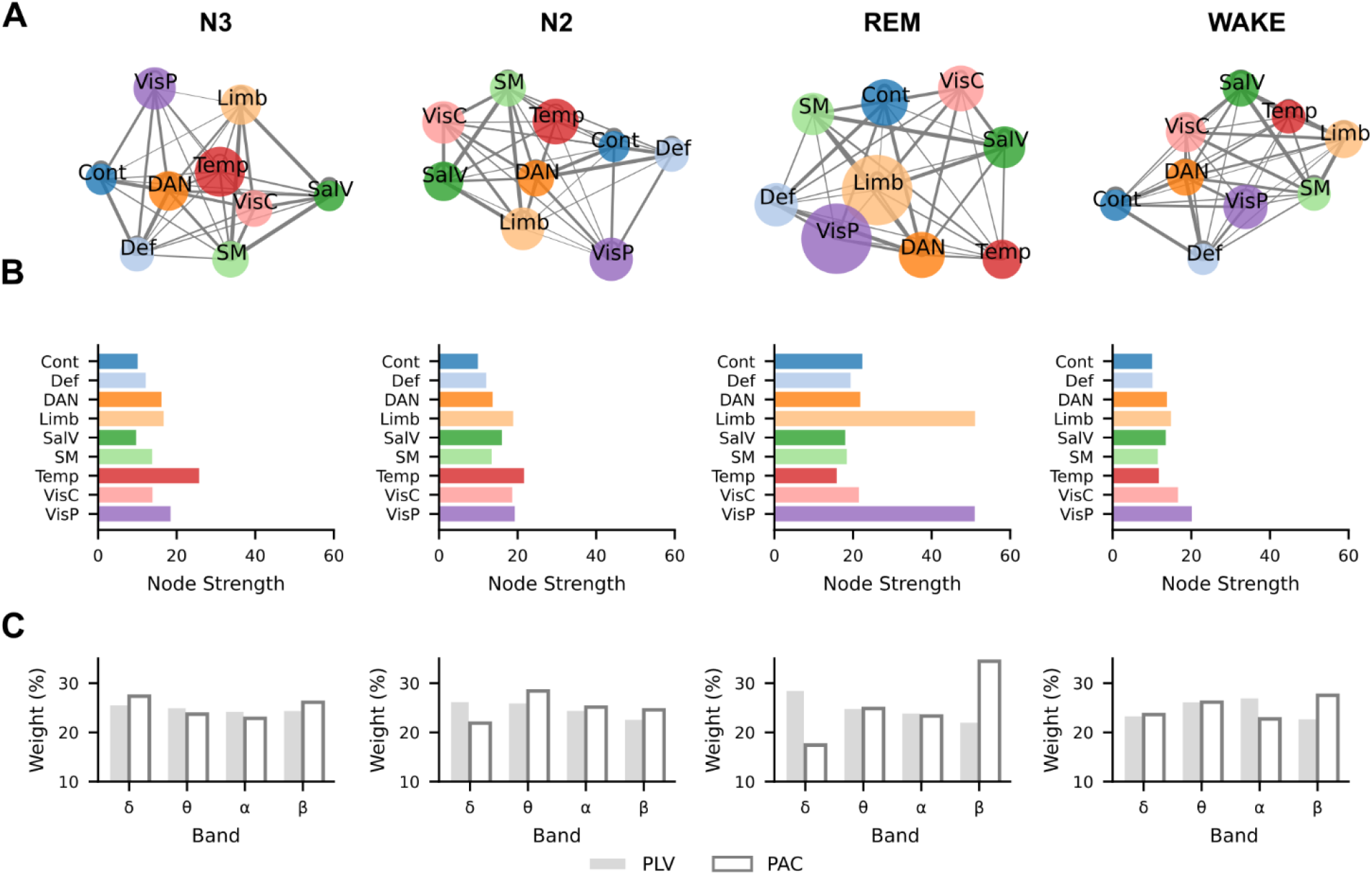
Joint representation of synchrony across functional systems. **(A)** Graph representations (spring layout) showing the subject-average scores of PLV and PAC for each vigilance state (n_N3_ = 38, n_N2_= 27, n_REM_ = 23, n_WAKE_ = 38). Node size is proportional to node strength (computed as s_i_ = ∑_j_ w_ij_, where wi_j_ is the edge weight), and edge thickness reflects the number of subjects exhibiting that connection. **(B)** Node strength of each functional system for each vigilance state. **(C)** PLS weights for PLV and PAC, shown separately for each vigilance state, indicating the relative contribution of each frequency band to the PLV-PAC covariation.

## Discussion

Our results reveal that large-scale human cortical dynamics reorganize across vigilance states, expressed through coordinated changes in phase synchronization and cross-frequency coupling. Together, these findings outline a multi-scale architecture in which slow rhythms dynamically gate faster activity, with distinct network signatures emerging across wakefulness, NREM, and REM sleep.

Inter-areal phase synchronization showed selective peaks in the δ, θ, σ, and low-β frequencies, highlighting their sensitivity to behavioural state. During NREM, θ and σ synchronization dominated across widespread regions, in agreement with the prominence of slow oscillations and spindle processes that structure memory reactivation during deep sleep (Diekelmann and Born, 2010; Klinzing et al., 2019). By contrast, REM exhibited heightened β and ripple-range coupling, consistent with fast, internally driven dynamics and coordinated high-frequency activity associated with hippocampo-cortical replay and dreaming (Boyce et al., 2016; Simor et al., 2018, 2019). These dissociations indicate that vigilance states impose principled constraints on the frequency channels supporting long-range interactions, shifting the balance between slow, spatially coherent coordination and faster, more selective interactions.

Phase-amplitude coupling further emphasized this architecture. During N3, δ-phase modulated broadband high-frequency activity, with spindle coupling providing an additional organizing axis (Bp et al., 2015; Latchoumane et al., 2017). As sleep lightened into N2, coupling expanded to include θ and σ phase influences on β and γ amplitudes, positioning N2 as an intermediate state with both slow oscillatory scaffolding and higher-frequency interactions. REM exhibited globally reduced PAC, consistent with attenuated hierarchical nesting despite preserved fast bursts associated (Niethard et al., 2016). Wakefulness showed θ-to-β coupling, characteristic of cortico-subcortical engagement during active cognitive processing (Canolty et al., 2006; Arnulfo et al., 2020).

We next asked how epileptogenic tissue is embedded within these dynamics. Electrodes in the epileptogenic zone (EZ) showed increases in δ and γ synchronization, consistent with a persistent imbalance in excitation– inhibition and hypersynchrony characteristic of epileptogenic networks (Wang et al., 2024; Burlando et al., 2026). Cross-frequency interactions were likewise elevated, with δ-to-β/γ/high-γ coupling particularly enhanced during N2 and wakefulness, supporting the notion that slow rhythms pathologically entrain fast microcircuit generators within the EZ (Jacobs et al., 2012; Frauscher et al., 2015b; Burlando et al., 2026).

REM did not exhibit PAC differences between EZ and non-EZ contacts, paralleling the reduced seizure probability during REM sleep (Ng and Pavlova, 2013; Amiri et al., 2016; Nobili et al., 2025). These observations suggest that REM-specific desynchronization and neuromodulatory configuration may actively suppress pathological slow–fast entrainment, reducing the capacity of EZ microcircuits to self-coordinate into seizure-promoting patterns (Frauscher et al., 2016; Nobili et al., 2025). To determine which large-scale systems scaffold these dynamics, we applied partial least squares linking PLV and PAC. NREM was dominated by temporal networks, with θ PAC contributions during N2 and δ coupling during N3, consistent with hippocampo-temporal replay mechanisms (Diekelmann and Born, 2010; Bp et al., 2015; Klinzing et al., 2019). Within systems-consolidation frameworks, NREM coordination is often described as a temporally nested interaction between slow oscillations, spindles, and ripple-range events that supports hippocampo-cortical communication and cortical reinstatement (Geva-Sagiv et al., 2023). From this perspective, a temporal-system hub may reflect the prominent role of temporo-hippocampal and temporo-association pathways in organizing replay and integration during NREM, even when the ultimate long-range target (*e*.*g*., prefrontal cortex) is distributed across multiple functional systems rather than expressed as a single dominant “frontal” node. In REM sleep, coupling reoriented toward visual and limbic circuits, dominated by δ-PLV and β-PAC, reflecting the integration of perceptual and affective processes that underlie dream generation. (Scarpelli et al., 2019). Notably, the strongest hub emerged in the peripheral/lateral visual system (VisP), rather than the central/medial visual system (VisC). In Schaefer-type parcellations (Schaefer et al., 2018), this “peripheral” subdivision maps preferentially onto more lateral/extra-striate visual territories (*i*.*e*., beyond primary calcarine cortex) and reflects supra-areal organization such as visual-field eccentricity. This bias toward higher-order visual regions is consistent with the phenomenology of REM dreaming, vivid, often emotionally salient, internally generated imagery, which is more likely to depend on associative visual processing in posterior cortex than on bottom-up encoding in early retinotopic cortex. These REM signatures align with seminal functional neuroimaging evidence showing preferential engagement of limbic and temporo-occipital cortices during REM, accompanied by relative prefrontal deactivation, a configuration that has been linked to the emotional salience, visual vividness, and reduced executive monitoring of dream experience (Maquet et al., 1996; Braun et al., 1997; Dang-Vu et al., 2010). Wakefulness exhibited a decentralized coupling architecture, indicative of dynamic reconfiguration across sensory and associative systems supporting adaptive cognition (Barttfeld et al., 2015; Mattar et al., 2015).

Together, these observations demonstrate that phase synchronization and PAC coupling offer complementary access to the structure of human sleep–wake dynamics. Although SEEG sampling is spatially sparse and determined by clinical necessity, the convergence of spectral, coupling, and systems findings suggests that similar motifs likely extend beyond the sampled network. In addition, statistical coupling does not by itself establish causal flow; directed and perturbational measures will be needed to determine how these motifs support computation or constrain ictogenesis. Longitudinal studies may further reveal whether these signatures evolve with learning or predict clinical outcome. Collectively, our results highlight principled constraints governing electrophysiological communication across sleep-wake states and show how pathological circuitry is embedded within and shaped by these dynamic regimes.

## References

Adamantidis, A. R., Gutierrez Herrera, C., and Gent, T. C. (2019). Oscillating circuitries in the sleeping brain.Nat Rev Neurosci20, 746–762. doi: 10.1038/s41583-019-0223-4

Amiri, M., Frauscher, B., and Gotman, J. (2016). Phase-Amplitude Coupling Is Elevated in Deep Sleep and in the Onset Zone of Focal Epileptic Seizures. Front Hum Neurosci10, 387. doi: 10.3389/fnhum.2016.00387

Arnulfo, G., Hirvonen, J., Nobili, L., Palva, S., and Palva, J. M. (2015). Phase and amplitude correlations in resting-state activity in human stereotactical EEG recordings. Neuroimage112, 114–127. doi: 10.1016/j.neuroimage.2015.02.031

Arnulfo, G., Wang, S. H., Myrov, V., Toselli, B., Hirvonen, J., Fato, M. M., et al. (2020). Long-range phase synchronization of high-frequency oscillations in human cortex. Nat Commun11, 5363. doi: 10.1038/s41467-020-18975-8

Barttfeld, P., Uhrig, L., Sitt, J. D., Sigman, M., Jarraya, B., and Dehaene, S. (2015). Signature of consciousness in the dynamics of resting-state brain activity. Proc Natl Acad Sci U S A112, 887–892. doi: 10.1073/pnas.1418031112

Bonnefond, M., Kastner, S., and Jensen, O. (2017). Communication between Brain Areas Based on Nested Oscillations. eNeuro4, ENEURO.0153-16.2017. doi: 10.1523/ENEURO.0153-16.2017

Boyce, R., Glasgow, S. D., Williams, S., and Adamantidis, A. (2016). Causal evidence for the role of REM sleep theta rhythm in contextual memory consolidation. Science352, 812–816. doi: 10.1126/science.aad5252

Bp, S., To, B., M, B., R, van der M., O, J., L, D., et al. (2015). Hierarchical nesting of slow oscillations, spindles and ripples in the human hippocampus during sleep. Nature neuroscience18. doi: 10.1038/nn.4119

Braun, A. R., Balkin, T. J., Wesenten, N. J., Carson, R. E., Varga, M., Baldwin, P., et al. (1997). Regional cerebral blood flow throughout the sleep-wake cycle. An H2(15)O PET study. Brain 120 (Pt 7), 1173–1197. doi: 10.1093/brain/120.7.1173

Burlando, G., Belforte, C., Siebenhühner, F., Tullio, L. D., Chiarella, L., Myrov, V., et al. (2026). Unstable slow oscillations couple with epileptogenic fast-rhythm bistability in sleep-related epilepsy: A stereoelectroencephalographic study. Epilepsia. doi: 10.1002/epi.70188

Cannon, J., McCarthy, M. M., Lee, S., Lee, J., Börgers, C., Whittington, M. A., et al. (2014). Neurosystems: brain rhythms and cognitive processing. Eur J Neurosci39, 705–719. doi: 10.1111/ejn.12453

Canolty, R. T., Edwards, E., Dalal, S. S., Soltani, M., Nagarajan, S. S., Kirsch, H. E., et al. (2006). High gamma power is phase-locked to theta oscillations in human neocortex. Science313, 1626–1628. doi: 10.1126/science.1128115

Canolty, R. T., and Knight, R. T. (2010). The functional role of cross-frequency coupling. Trends Cogn Sci14, 506–515. doi: 10.1016/j.tics.2010.09.001

Chiarion, G., Sparacino, L., Antonacci, Y., Faes, L., and Mesin, L. (2023). Connectivity Analysis in EEG Data: A Tutorial Review of the State of the Art and Emerging Trends. Bioengineering (Basel)10, 372. doi: 10.3390/bioengineering10030372

Corsi-Cabrera, M., Guevara, M. A., Arce, C., and Ramos, J. (1996). Inter and intrahemispheric EEG correlation as a function of sleep cycles. Prog Neuropsychopharmacol Biol Psychiatry20, 387–405. doi: 10.1016/0278-5846(96)00004-8

Corsi-Cabrera, M., Muñoz-Torres, Z., del Río-Portilla, Y., and Guevara, M. A. (2006). Power and coherent oscillations distinguish REM sleep, stage 1 and wakefulness. Int J Psychophysiol60, 59–66. doi: 10.1016/j.ijpsycho.2005.05.004

Cox, R., Rüber, T., Staresina, B. P., and Fell, J. (2020). Phase-based coordination of hippocampal and neocortical oscillations during human sleep. Commun Biol3, 1–11. doi: 10.1038/s42003-020-0913-5

Dang-Vu, T. T., Schabus, M., Desseilles, M., Sterpenich, V., Bonjean, M., and Maquet, P. (2010). Functional neuroimaging insights into the physiology of human sleep. Sleep33, 1589–1603. doi: 10.1093/sleep/33.12.1589

Diekelmann, S., and Born, J. (2010). The memory function of sleep. Nat Rev Neurosci11, 114–126. doi: 10.1038/nrn2762

Engel, A. K., and Fries, P. (2010). Beta-band oscillations--signalling the status quo? Curr Opin Neurobiol20, 156–165. doi: 10.1016/j.conb.2010.02.015

Fischl, B. (2012). FreeSurfer. NeuroImage62, 774–781. doi: 10.1016/j.neuroimage.2012.01.021

Frauscher, B., von Ellenrieder, N., Dubeau, F., and Gotman, J. (2016). EEG desynchronization during phasic REM sleep suppresses interictal epileptic activity in humans. Epilepsia57, 879–888. doi: 10.1111/epi.13389

Frauscher, B., von Ellenrieder, N., Ferrari-Marinho, T., Avoli, M., Dubeau, F., and Gotman, J. (2015a). Facilitation of epileptic activity during sleep is mediated by high amplitude slow waves. Brain138, 1629–1641. doi: 10.1093/brain/awv073

Frauscher, B., von Ellenrieder, N., Ferrari-Marinho, T., Avoli, M., Dubeau, F., and Gotman, J. (2015b). Facilitation of epileptic activity during sleep is mediated by high amplitude slow waves. Brain138, 1629–1641. doi: 10.1093/brain/awv073

Fries, P. (2005). A mechanism for cognitive dynamics: neuronal communication through neuronal coherence.Trends Cogn Sci9, 474–480. doi: 10.1016/j.tics.2005.08.011

Genovese, C. R., Lazar, N. A., and Nichols, T. (2002). Thresholding of statistical maps in functional neuroimaging using the false discovery rate. Neuroimage15, 870–878. doi: 10.1006/nimg.2001.1037

Geva-Sagiv, M., Mankin, E. A., Eliashiv, D., Epstein, S., Cherry, N., Kalender, G., et al. (2023). Augmenting hippocampal–prefrontal neuronal synchrony during sleep enhances memory consolidation in humans. Nat Neurosci26, 1100–1110. doi: 10.1038/s41593-023-01324-5

Geva-Sagiv, M., and Nir, Y. (2019). Local Sleep Oscillations: Implications for Memory Consolidation. Front Neurosci13, 813. doi: 10.3389/fnins.2019.00813

Jacobs, J., Staba, R., Asano, E., Otsubo, H., Wu, J. Y., Zijlmans, M., et al. (2012). High-frequency oscillations (HFOs) in clinical epilepsy. Prog Neurobiol98, 302–315. doi: 10.1016/j.pneurobio.2012.03.001

Klinzing, J. G., Niethard, N., and Born, J. (2019). Mechanisms of systems memory consolidation during sleep.Nat Neurosci22, 1598–1610. doi: 10.1038/s41593-019-0467-3

Latchoumane, C.-F. V., Ngo, H.-V. V., Born, J., and Shin, H.-S. (2017). Thalamic Spindles Promote Memory Formation during Sleep through Triple Phase-Locking of Cortical, Thalamic, and Hippocampal Rhythms. Neuron95, 424–435.e6. doi: 10.1016/j.neuron.2017.06.025

Lehnertz, K., Bialonski, S., Horstmann, M.-T., Krug, D., Rothkegel, A., Staniek, M., et al. (2009). Synchronization phenomena in human epileptic brain networks. Journal of Neuroscience Methods183, 42–48. doi: 10.1016/j.jneumeth.2009.05.015

Maquet, P., Péters, J., Aerts, J., Delfiore, G., Degueldre, C., Luxen, A., et al. (1996). Functional neuroanatomy of human rapid-eye-movement sleep and dreaming. Nature383, 163–166. doi: 10.1038/383163a0

Mattar, M. G., Cole, M. W., Thompson-Schill, S. L., and Bassett, D. S. (2015). A Functional Cartography of Cognitive Systems. PLoS Comput Biol11, e1004533. doi: 10.1371/journal.pcbi.1004533

Muñoz-Torres, Z., Corsi-Cabrera, M., Velasco, F., and Velasco, A. L. (2023). Amygdala and hippocampus dialogue with neocortex during human sleep and wakefulness. Sleep46, zsac224. doi: 10.1093/sleep/zsac224

Myrov, V., Siebenhühner, F., Wang, S. H., Arnulfo, G., Juvonen, J., Roascio, M., et al. (2026). CROCOpy - A Python toolbox for the analysis of CRitical Oscillations and COnnectivity. 2026.02.17.706438. doi: 10.64898/2026.02.17.706438

Narizzano, M., Arnulfo, G., Ricci, S., Toselli, B., Tisdall, M., Canessa, A., et al. (2017). SEEG assistant: a 3DSlicer extension to support epilepsy surgery. BMC Bioinformatics18, 124. doi: 10.1186/s12859-017-1545-8

Ng, M., and Pavlova, M. (2013). Why are seizures rare in rapid eye movement sleep? Review of the frequency of seizures in different sleep stages. Epilepsy Res Treat 2013, 932790. doi: 10.1155/2013/932790

Niethard, N., Hasegawa, M., Itokazu, T., Oyanedel, C. N., Born, J., and Sato, T. R. (2016). Sleep-Stage-Specific Regulation of Cortical Excitation and Inhibition. Current Biology26, 2739–2749. doi: 10.1016/j.cub.2016.08.035

Nobili, L., Cordani, R., Arnaldi, D., Mattioli, P., Veneruso, M., and Ng, M. (2025). Rapid eye movement sleep and epilepsy: exploring interactions and therapeutic prospects. Journal of Sleep Research34, e14251. doi: 10.1111/jsr.14251

Nobili, L., De Gennaro, L., Proserpio, P., Moroni, F., Sarasso, S., Pigorini, A., et al. (2012). “Chapter 13 - Local aspects of sleep: Observations from intracerebral recordings in humans,” in Progress in Brain Research, eds. A. Kalsbeek, M. Merrow, T. Roenneberg, and R. G. Foster (Elsevier), 219–232. doi: 10.1016/B978-0-444-59427-3.00013-7

Palmigiano, A., Geisel, T., Wolf, F., and Battaglia, D. (2017). Flexible information routing by transient synchrony. Nat Neurosci20, 1014–1022. doi: 10.1038/nn.4569

Palva, J. M., Wang, S. H., Palva, S., Zhigalov, A., Monto, S., Brookes, M. J., et al. (2018). Ghost interactions in MEG/EEG source space: A note of caution on inter-areal coupling measures. NeuroImage173, 632– 643. doi: 10.1016/j.neuroimage.2018.02.032

Roliz, A. H., and Kothare, S. (2022). The Interaction Between Sleep and Epilepsy. Curr Neurol Neurosci Rep22, 551–563. doi: 10.1007/s11910-022-01219-1

Scarpelli, S., Bartolacci, C., D’Atri, A., Gorgoni, M., and De Gennaro, L. (2019). The Functional Role of Dreaming in Emotional Processes. Front. Psychol.10. doi: 10.3389/fpsyg.2019.00459

Schaefer, A., Kong, R., Gordon, E. M., Laumann, T. O., Zuo, X.-N., Holmes, A. J., et al. (2018). Local-Global Parcellation of the Human Cerebral Cortex from Intrinsic Functional Connectivity MRI. Cereb Cortex28, 3095–3114. doi: 10.1093/cercor/bhx179

Schmidt, H., Petkov, G., Richardson, M. P., and Terry, J. R. (2014). Dynamics on Networks: The Role of Local Dynamics and Global Networks on the Emergence of Hypersynchronous Neural Activity. PLOS Computational Biology10, e1003947. doi: 10.1371/journal.pcbi.1003947

Schreiner, T., Griffiths, B. J., Kutlu, M., Vollmar, C., Kaufmann, E., Quach, S., et al. (2024). Spindle-locked ripples mediate memory reactivation during human NREM sleep. Nat Commun15, 5249. doi: 10.1038/s41467-024-49572-8

Sheybani, L., Frauscher, B., Bernard, C., and Walker, M. C. (2025). Mechanistic insights into the interaction between epilepsy and sleep. Nature Reviews Neurology, 1–16. Available at: https://www.nature.com/articles/s41582-025-01064-z (Accessed October 8, 2025).

Simor, P., Gombos, F., Blaskovich, B., and Bódizs, R. (2018). Long-range alpha and beta and short-range gamma EEG synchronization distinguishes phasic and tonic REM periods. Sleep41. doi: 10.1093/sleep/zsx210

Simor, P., van Der Wijk, G., Gombos, F., and Kovács, I. (2019). The paradox of rapid eye movement sleep in the light of oscillatory activity and cortical synchronization during phasic and tonic microstates. NeuroImage202, 116066. doi: 10.1016/j.neuroimage.2019.116066

Singer, W. (1999). Neuronal synchrony: a versatile code for the definition of relations? Neuron24, 49–65, 111–125. doi: 10.1016/s0896-6273(00)80821-1

Staresina, B. P., Niediek, J., Borger, V., Surges, R., and Mormann, F. (2023). How coupled slow oscillations, spindles and ripples control neuronal processing and communication during human sleep. Neuroscience. doi: 10.1101/2023.01.08.523138

Tononi, G., Boly, M., Massimini, M., and Koch, C. (2016). Integrated information theory: from consciousness to its physical substrate. Nat Rev Neurosci17, 450–461. doi: 10.1038/nrn.2016.44

Varela, F., Lachaux, J. P., Rodriguez, E., and Martinerie, J. (2001). The brainweb: phase synchronization and large-scale integration. Nat Rev Neurosci2, 229–239. doi: 10.1038/35067550

Vyazovskiy, V. V., Olcese, U., Hanlon, E. C., Nir, Y., Cirelli, C., and Tononi, G. (2011). Local sleep in awake rats. Nature472, 443–447. doi: 10.1038/nature10009

Wang, S. H., Arnulfo, G., Nobili, L., Myrov, V., Ferrari, P., Ciuciu, P., et al. (2024). Neuronal synchrony and critical bistability: Mechanistic biomarkers for localizing the epileptogenic network. Epilepsia65, 2041–2053. doi: 10.1111/epi.17996

Yeo, B. T. T., Krienen, F. M., Sepulcre, J., Sabuncu, M. R., Lashkari, D., Hollinshead, M., et al. (2011). The organization of the human cerebral cortex estimated by intrinsic functional connectivity. J Neurophysiol106, 1125–1165. doi: 10.1152/jn.00338.2011

