## Supplementary Material for "Frequency modulations of cortical synchronization in human cortex during wakefulness and sleep"

Canu et al.

### Supplementary Figures

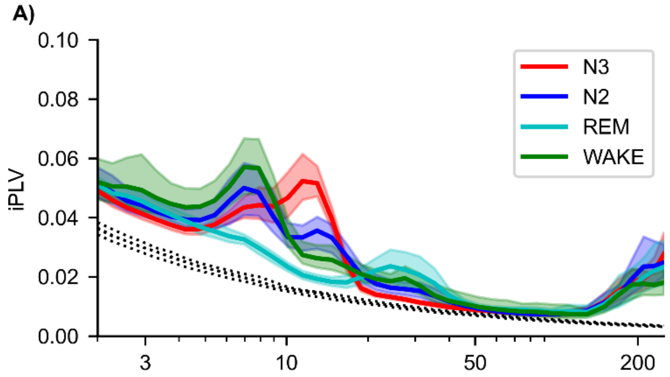


***Figure S1. (A)*** *iPLV spectra for N3 (red), N2 (blue), REM (cyan), and wake (green) conditions across subjects in nEZ. The shaded areas represent the 2.5th to 97.5th percentile bootstrap (N=1000) confidence limits around the mean (thick lines).*

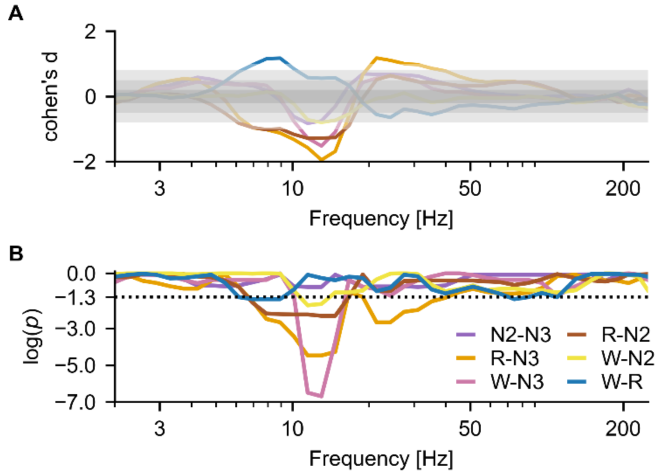


***Figure S2. (A)*** *Effect size computed with Cohen's d for the comparison of PLV pairwise for vigilance states. Coloured areas represent the value range for small (|d|<0.2, dark grey), medium (0.2<|d|<0.8, light grey), and large (|d|> 0.8, dark grey) effect sizes. (B) p-values from the Wilcoxon signed-rank test for the comparison of PLV pairwise for vigilance states. The black dashed line denotes the significance threshold (p = 0.05).*
